# miRNAgFree: prediction and profiling of novel microRNAs without genome assembly

**DOI:** 10.1101/193094

**Authors:** EL Aparicio, A Rueda, B. Fromm, C Gómez-Martín, R Lebrón, JL Oliver, JA Marchal, M Kotsyfakis, M Hackenberg

**Affiliations:** Bioinformatics and Epigenomics Lab, Genetics Dpt. Faculty of Science, University of Granada, Granada, Spain; Biotechnology Institute, University of Granada, Av. del Conocimiento, Granada; Genomics England, Charterhouse Square, London EC1M 6BQ, UK; Department of Tumor Biology, Institute for Cancer Research, Oslo University Hospital, Oslo, Norway; Department of Human Anatomy and Embryology, Institute of Biopathology and Regenerative Medicine, University of Granada, Granada, Spain; Biology Center of the Czech Academy of Sciences, Budweis, Czech Republic

**Keywords:** Bioinformatics, reference genome, prediction, de novo, genome

## Abstract

The prediction of novel miRNA genes generally requires the availability of genome sequences in order to assess important properties such as the characteristic hairpin-shaped secondary structure. However, although the sequencing costs have decreased over the last years, still many important species lack an assembled genome of certain quality. We implemented an algorithm which for the first time exploits characteristic biogenesis features like the 5’ homogeneity that can be assessed without genome sequences. We used a phylogenetically broad spectrum of well annotated animal genomes for benchmarking. We found that between 90-100% of the most expressed miRNA candidates (top quartile) corresponded to known miRNA sequences.

## 1 Introduction

MicroRNAs (miRNAs) have important roles in many biological processes (Bushati and Cohen, 2007) and they possess a huge potential to become prominent biomarkers as they can be detected in nearly every bodily fluid (Cortez et al., 2011). The miRNA expression profiling can be routinely carried out by means of micro-arrays or next-generation-sequencing if the mature and pre-miRNA sequences are available. Therefore the determination of the miRNA reference sequences is an essential task. Generally, genome sequences are required and many different programs are available for the prediction of novel miRNAs like miRanalyzer and miRDeep2 (Hackenberg et al., 2011; Friedländer et al., 2012) or miRCandRef which works with unassembled reads (Fromm et al., 2013). There are also some programs available for the prediction without genome like miRMiner (Wheeler et al., 2009) which is based on homology and, miRPlex (Mapleson et al., 2013) and miReader (Jha et al., 2013) based on machine learning using different duplex features.

Here we present a novel miRNA prediction approach based on biogenesis features, such as the known 5’ homogeneity, and duplex features like mean free energy which do not require a genome assembly to be assessed. We found that, in general, biogenesis related parameters are far more discriminative than duplex related structural parameters. We observed that a high percentage of the top expressed miRNA candidates in animals correctly match actual guide sequences while the prediction without genome in plants seems to be more complex, leading to much lower specificities. Our approach outperforms previous similar attempts because microRNA biogenesis features were taken into account. We benchmarked miRNAgFree using a set of species with high quality genomes (including *H.sapiens, mus musculus, C.elegans, D.Rerio)* using publicly available datasets. When measuring the specificity on the guide strand of the microRNA we obtained over 90% accuracy for the most expressed quartile of duplexes.

## 2 Main features and implementation

miRNAgFree is a piece of software that allows for mature microRNA prediction using sRNAseq/miRNAseq/sncRNAseq without needing a genome or miRNA sequence libraries. This tool is therefore ideal for non-model species with genomes yet to be sequenced or for those lacking an appropriate quality. The software uses the sRNAbench preprocessing and therefore accepts several input files:

- Adapter trimming can be performed and miRNAgFree accepts *fastq*, *fastq.gz*, read count and *fasta* input format
- A preprocessing filtering step can be included. This is useful in a number of scenarios: for example to remove unwanted ribosomal sequences or if there are some already described microRNAs that should be eliminated from the analysis. Reads mapping to the provided libraries will not be considered for downstream analysis.
- Lax parameter settings (more sensitive) and strict settings (more specific) are provided.

### 2.1 Implementation steps

The general workflow consists in i) preprocessing of the input reads, ii) read filtering (optional if the user provides a filter library like ribosomal RNA), iii) clustering of the reads, iv) calculation of duplexes with RNAcofold, v) detection of microRNA-like duplexes and vi) output layer including visualization of the detected miRNA duplexes.

#### 2.1.1 Cluster method

The mature microRNAs are normally represented at the read level by the canonical sequence (i.e. the one in miRBase) and its isomiR sequences. Therefore, in order not to predict a microRNA several times due to duplexes formed by its isomiR sequences, we first cluster together all reads. Briefly the algorithm performs the following steps on a sorted read list (descending order)

1. Open a cluster with the most abundant read as dominant read
2. Take the most abundant read as reference and align all other reads against it. The reads can align with a 3 nt overhang at the 5’ end allowing by default 1 mismatch (suppl. figure 1a). The last nucleotides are ignored as those might be NTAs (non-templated additions)
3. Remove all assigned reads so they are not considered in other clusters
4. Repeat steps 1-3 until no reads are left.

**Fig 1.**
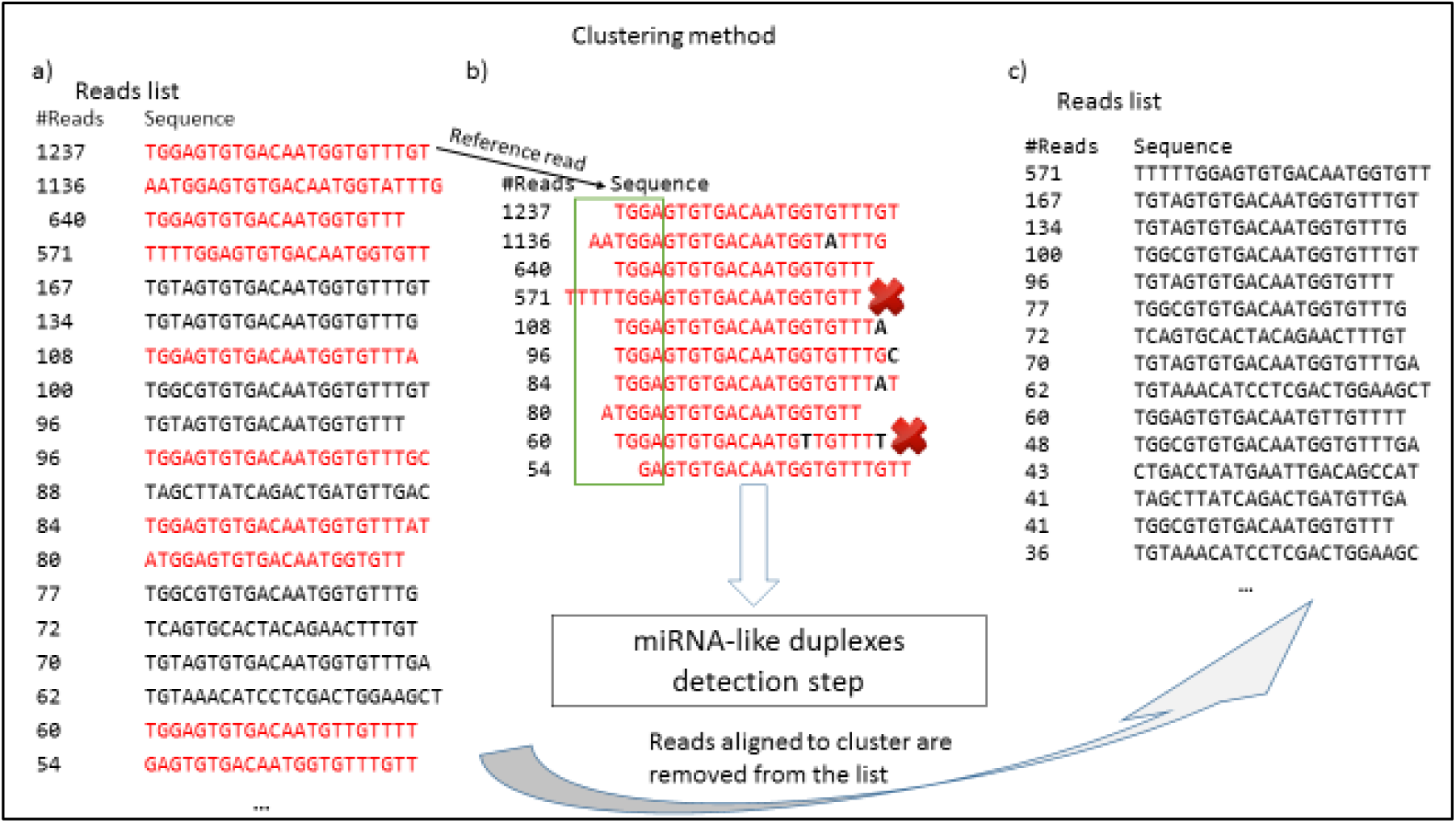
Iterative clustering method. a) The most expressed read opens the cluster and all reads are aligned to this sequence. b) All reads that map to the reference are part of the cluster. Reads need to start within a 3nt window around the 5’ end of the dominant read (green rectangle) and 1 mismatch (nucleotides in black) is allowed by default (user parameter). c) All assigned reads are removed from the input list and the iterative steps start again at point a).

#### 2.1.2 Detection of miRNA duplexes

The detection of novel microRNAs consists of two steps: i) selection of clusters that resemble those formed by real guide microRNAs and their isomiRs and ii) the assignment of the most likely passenger sequence.

1. Sort clusters by total expression value in descending order.
2. Pick the most expressed cluster and remove it from the list and evaluate its microRNA potential based on several criteria like the 5’ homogeneity or the ratio of the most expressed read to the read count sum of the cluster. The same thresholds as in (Barturen et al., 2014) were used. The user can choose between lax and strict settings for both animals and plants.
3. For all putative cluster pairs, calculate all duplexes for the (M) most expressed reads in cluster (i) vs the (N) most expressed clusters of cluster (j) and remove all that do not have perfect 2 nucleotide 3’ Drosha/Dicer overhangs (strict) or at least 1-3 nt overhang (lax mode)
4. Sort the duplexes by energy ratio (mean free energy divided by the sequence length) and assign the energetically most favorable read and its cluster to the guide cluster.

**Fig. 2.**
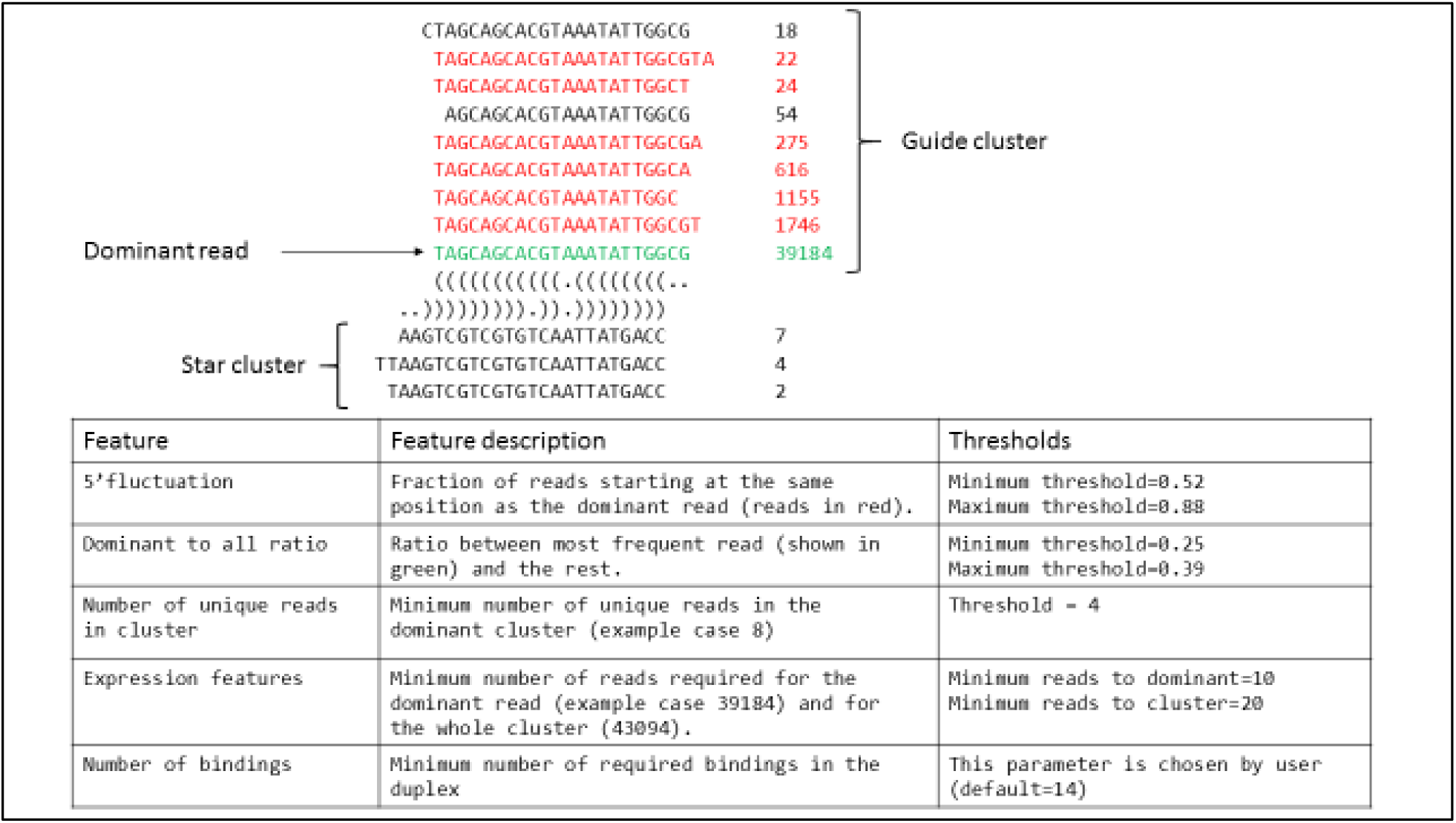
Parameters and thresholds used in the miRNA-like selection

## 3 Results

To show the usefulness of our approach we used two strategies: i) assess the number of correct predictions by means of well annotated genomes and ii) show that species with an unfinished genome assembly (*Fasciola hepatica*) very likely have incomplete miRNA complements. We found that the percentage of correct guide sequences ranges from 51% (C. elegans) to 91% (D. melanogaster) in animals but are generally lower in plants. The fraction of correct guide sequences increases with the threshold number of bindings, whereas the total number of predictions drops strongly, and hence the sensitivity. The prediction quality seems to be notably affected by the quality of the sequencing data. For the M. musculus data we observed a specificity increase from 50.8% to 85.2% when filtering out all reads that have a single position with lower phred score than 20. Even though we prioritized specificity to increase confidence in the yielded prediction, we still found that sensitivity ranged between 51% (most strict set of parameters) and 81% for the least strict settings (see suppl. Table 1).

Further, we found that prediction quality increases with expression value, i.e. the top 14 guide sequences in mouse and the top 44 in zebrafish are all correct (see column c in Table 1). Finally, using publically available data for *F. hepatica* (SRR1825354) we detected a member of the let-7 family which was not reported in the most recent complement using the genome assembly (Fromm et al., 2017) either due to assembly quality or structural properties which prevented its prediction (see supplementary figure 1).

**Table 1.** Benchmark of miRNAgFree using well annotated species.

| Species | b | N | p (%) | p25 (%) | c | SRA accession |
| --- | --- | --- | --- | --- | --- | --- |
| *Homo sapiens* | 16 | 79 | 82.3 | 95.7 | 16 | SRR1563015 |
| *Homo sapiens* | 18 | 24 | 91.7 | 100 | 22 | SRR1563015 |
| *Mus musculus* | 16 | 61 | 85.2 | 93.3 | 14 | SRR1734811 |
| *Mus musculus* * | 16 | 118 | 50.8 | 90 | 14 | SRR1734811 |
| *Danio rerio* | 16 | 170 | 75.9 | 100 | 44 | SRR3953259 |
| *D. melanogaster* | 16 | 33 | 90.9 | 100 | 29 | SRR1287661 |
| *C. elegans* | 16 | 183 | 51.4 | 91.3 | 40 | ERR562747 |
| *A. thaliana* | 16 | 63 | 30.1 | 37.5 | 2 | SRR5031522 |

b is the applied duplex bindings threshold, N the total number of predicted miRNAs, p is the proportion of correct guide sequences (annotated in miRBase 21), p25 is the same as p but just considering the top quartile of predictions, c is the number of consecutive correct predictions among the top expressed ones. \* indicates that no quality filter was applied to the raw fastq file.

In summary, we tested different duplex related structural features as used in prior approaches, but we found that biogenesis features and specially the 5’ homogeneity outperform those clearly. Given that the top expressed guide miRNAs are generally correct, miRNAgFree can be an important tool for miRNA research in non-model species.

## Supplementary material

**Supplementary table 1.** Specificity and sensitivity values obtained for the *Homo sapiens* dataset

| Mode used | Maximum length allowed for candidate microRNA | Number of mismatches allowed in the clustering step | Sensitivity | Specificity |
| --- | --- | --- | --- | --- |
| Strict mode (same as Table 1) | 23 | 1 | 0.5112359551 | 0.79 |
| Lax mode (keeping the rest as in Table 1) | 23 | 1 | 0.6853932584 | 0.6338582677 |
| Lax mode | 24 | 1 | 0.7430167598 | 0.5906040268 |
| Lax mode | 24 | 0 | 0.808988764 | 0.7879464286 |

**Suppl. Fig 1.**
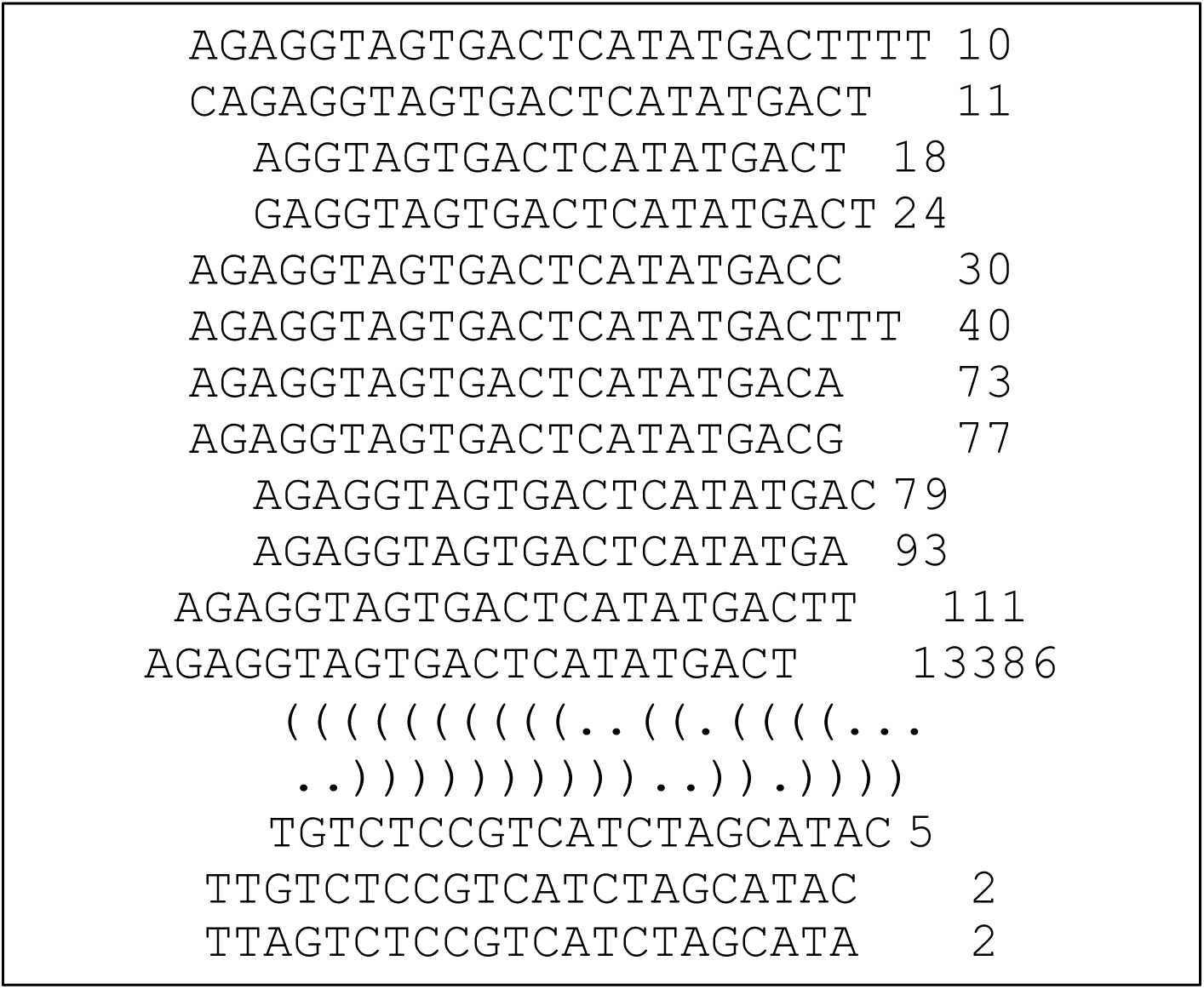
A real example from let-7 in *F. hepatica* predicted using the publically available dataset *SRR1825354*.

